# Epigenetic and 3D Genome Changes Drive Primary Trastuzumab Resistance in HER2+ Breast Cancer

**DOI:** 10.64898/2026.04.24.720531

**Authors:** Yijia Hua, Zhengxing Zhou, Nan Jin, Wei Li, Yongmei Yin, Ningjun Duan

**Author notes:** Corresponding to Yongmei Yin and Ningjun Duan. These authors contributed equally to the manuscript (Yijia Hua, Zhengxing Zhou).

## Abstract

Primary trastuzumab resistance remains a major challenge in the treatment of HER2-positive breast cancer, with limited therapeutic options highlighting the need for deeper mechanistic insight. Here, integrative analysis of epigenomic and three-dimensional chromatin architecture reveals widespread remodeling of the regulatory landscape in resistant cells. Alterations in promoter-associated histone modifications—particularly H3K4me3 and H3K27me3—together with changes in active enhancer activity and promoter–enhancer interactions, appear to be key drivers of transcriptional reprogramming underlying resistance. As an illustrative example, SGK1 exhibits epigenetic activation in resistant cells, characterized by increased promoter H3K4me3 and enhanced chromatin interactions, consistent with its role in supporting cell survival, proliferation and metastasis. Overall, our findings highlight epigenetic and chromatin structural alterations as central mechanisms of primary trastuzumab resistance, providing a framework for novel therapeutic strategy development.

## Introduction

Breast cancer is currently the most prevalent malignancy affecting women worldwide, with nearly a quarter of all cases being HER2-positive subtypes^1^. Although CDK4/6 inhibitors, tyrosine kinase inhibitors, and antibody-drug conjugates (ADCs) are increasingly used in later-stage treatment of HER2-positive breast cancer, trastuzumab remains the primary choice for early-stage treatment (neoadjuvant and adjuvant therapies)^2^. Nevertheless, a significant challenge persists in the treatment showing poor or no response to trastuzumab^3^. Several theories have explained the mechanisms of primary trastuzumab resistance, including, but not limited to, HER2 amplification and mutation, as well as the abnormal activation of downstream or bypass signaling pathways^4,5^. However, proposed solutions targeting these mechanisms have not been as effective as expected in clinical applications. All these still highlight the importance of further investigating the mechanisms of primary trastuzumab resistance.

The carcinogenesis of breast cancer involves a combination of complex genetic and epigenetic alterations that collectively drive the transformation of normal ductal epithelial cells into carcinoma cells^6^. While genetic alterations such as mutations, amplifications, deletions, and rearrangements result in permanent changes in gene function and expression, epigenetic alterations in DNA and histone modifications, as well as 3D chromatin structure, provide a more dynamic means of gene expression control for both oncogenes and tumor suppressor genes^7–10^. Among these, histone modifications at promoters, particularly tri-methylation of H3K4 and H3K27, are known to directly influence the binding of RNA polymerase II to promoter regions, thereby regulating the initiation of gene transcription^11,12^. Simultaneously, acylated H3K27-marked active enhancers can aid in recruiting RNA polymerase II to both nearby and distant promoters by forming chromatin loops with the cohesin complex^13–15^. All these mechanisms function collaboratively to regulate the transcription of genes during the progression of cancer. Although years of research have demonstrated the roles of epigenetic alterations throughout the entire progression of breast cancer, from early carcinogenesis to metastasis, our understanding of several key processes, including the development of drug resistance, remains limited.

In this study, we delineate the epigenetic and chromatin landscape of primary trastuzumab-resistant and -sensitive cells to uncover mechanisms underlying resistance. By integrating histone modification profiles, chromatin interactions, and transcriptomic data, we identify substantial epigenomic divergence between the two states. Notably, alternative promoter H3K4me3 enrichment, together with active enhancer activity and strengthened promoter–enhancer interactions, appears to drive transcriptional reprogramming. Among the epigenetically regulated genes, serum/glucocorticoid-regulated kinase 1 (SGK1) was upregulated in the intrinsically resistant cell model and may contribute to cellular phenotypes associated with tumor progression and trastuzumab resistance.

## Results

### Variant promoter and enhancer histone modifications

To characterize the epigenetic landscapes of trastuzumab-resistant and trastuzumab-sensitive cells, we used CUT&Tag to profile the genome-wide distribution of H3K27me3, H3K4me3, and H3K27ac. We identified 11,801 H3K27me3 peaks in JIMT1 cells and 11,644 in SKBR3 cells, including 5,970 and 5,831 cell line-specific peaks, respectively, and 5,813 peaks shared between the two cell lines. Cell line-specific peaks were distributed across promoters, introns, exons, and distal regions (Figure 1a–b). We also identified 8,939 H3K4me3 peaks in JIMT1 cells and 8,936 in SKBR3 cells, of which 1,840 and 1,837 were cell line-specific, respectively. These peaks were predominantly located at promoters (Figure 1c–d).

**Figure 1.**
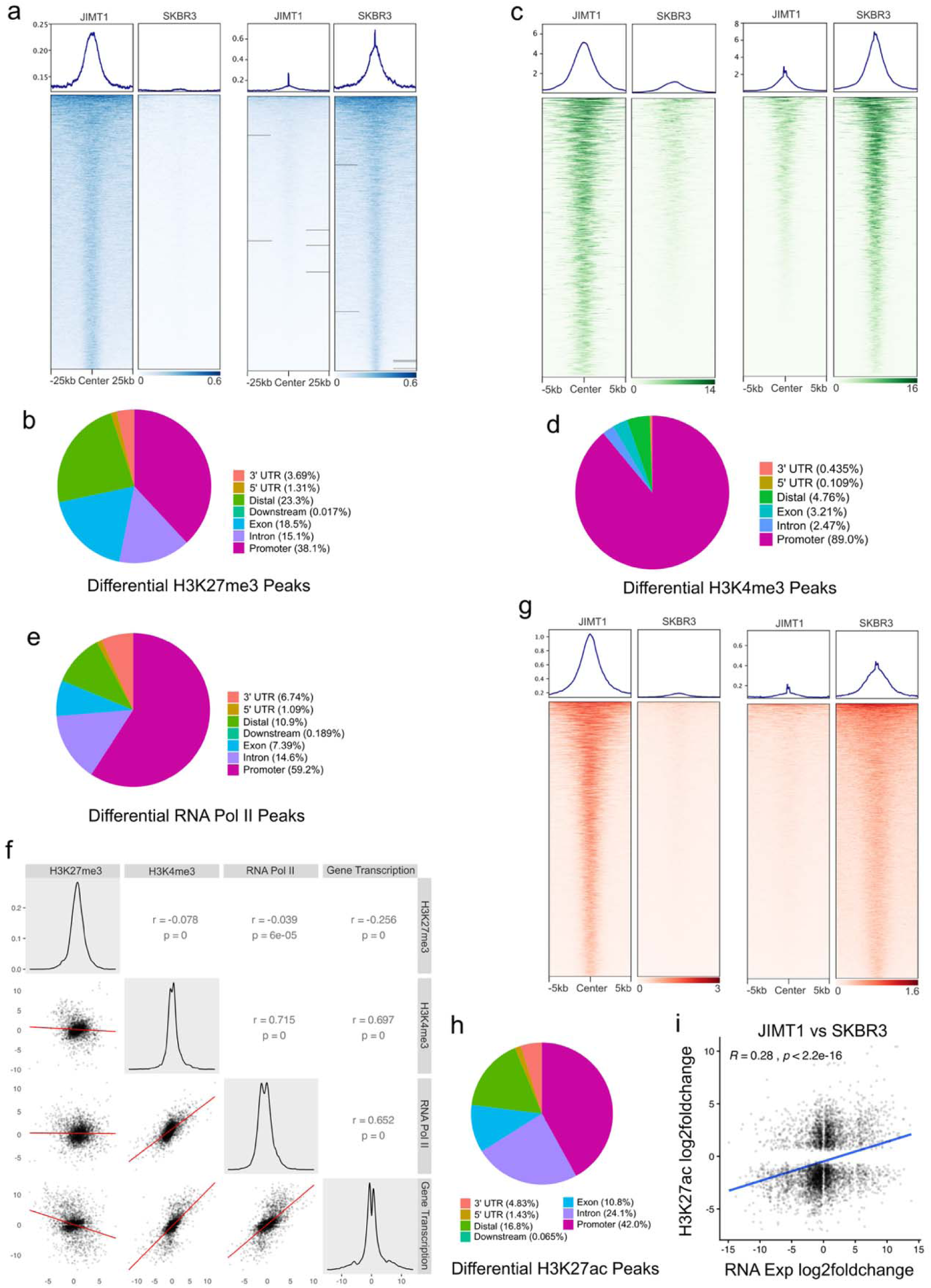
Altered histone modification between trastuzumab-sensitive and resistant cells. **a**, Differential H3K27me3 peaks in JIMT1 and SKBR3 cells. **b**, Distributions of differential H3K27me3 peaks in JIMT1 and SKBR3 cells. **c**, Differential H3K4me3 peaks in JIMT1 and SKBR3 cells. **d**, Distributions of differential H3K4me3 peaks in JIMT1 and SKBR3 cells. **e**, Distributions of differential RNA polymerase II peaks in JIMT1 and SKBR3 cells. **f**, Correlation among differential promoter H3K27me3, H3K4me3, RNA polymerase II enrichments and gene transcription. **g**, Differential H3K27ac peaks in JIMT1 and SKBR3 cells. **h**, Distributions of differential H3K27ac peaks in JIMT1 and SKBR3 cells. **i**, Pearson correlation between RNA expression and enhancer H3K27ac differences in JIMT1 and SKBR3 cells.

Because promoter-associated H3K27me3 and H3K4me3 are linked to transcriptional regulation, we also profiled RNA polymerase II using CUT&Tag. RNA polymerase II enrichment was predominantly detected at promoter regions in both cell lines (Figure 1e). We then examined the relationships among promoter histone modifications, RNA polymerase II occupancy, and gene expression. H3K4me3 enrichment was strongly positively correlated with both RNA polymerase II occupancy and transcriptional activity, whereas H3K27me3 showed a weak negative correlation. Promoter H3K27me3 levels were relatively stable across genes in both cell lines, while H3K4me3 and RNA polymerase II enrichment closely tracked gene expression. Highly expressed genes were associated with greater H3K4me3 enrichment and RNA polymerase II occupancy, whereas genes with stable expression showed corresponding stable patterns (Figure 1f). These observations suggest that promoter H3K4me3 enrichment is more closely associated with transcriptional activity than H3K27me3 in these cells.

The co-occurrence of H3K27me3 and H3K4me3 is a characteristic feature of bivalent promoters, which are associated with poised transcriptional states. Compared with SKBR3 cells, JIMT1 cells exhibited a gain of promoter H3K4me3 in more than 50% of highly expressed genes, a loss of H3K27me3 in about 5%, and concurrent H3K4me3 gain and H3K27me3 loss in nearly 22% (Figure S1a). Genes associated with activated bivalent promoters in JIMT1 cells were enriched in pathways and processes implicated in treatment resistance, including epithelial–mesenchymal transition, KRAS signaling, and TNF-α signaling (Figure S1b). These findings suggest that changes at bivalent promoters may be associated with the expression of resistance-related genes.

We next examined H3K27ac, a histone modification associated with active regulatory elements. CUT&Tag identified 17,824 H3K27ac peaks in JIMT1 cells and 18,316 in SKBR3 cells, including 9,065 and 9,667 peaks specific to the respective cell lines (Figure 1g). These peaks were predominantly distributed across promoters, introns, exons, and distal regions (Figure 1h). We identified H3K27ac-enriched regions outside promoters as putative enhancers. Although the total numbers of enhancers were comparable between JIMT1 and SKBR3 cells (Figure S1c), only 2,139 were shared, whereas 5,436 and 5,580 were specific to JIMT1 and SKBR3 cells, respectively (Figure S1d), indicating substantial differences in their enhancer landscapes.

In addition, gene expression differences generally correlated positively with H3K27ac levels at nearby enhancers within ±1 Mb in JIMT1 and SKBR3 cells (Figure 1i). Highly expressed genes were associated with higher enhancer H3K27ac, whereas lowly expressed genes were associated with lower enhancer H3K27ac (Figure S1e). These findings support an association between enhancer activity and transcriptional output.

### Differential genomic contacts and promoter-enhancer loops

Three-dimensional (3D) genome architecture reflects the folding of chromatin within the nucleus and contributes to the regulation of gene transcription. To investigate differences in genome architecture between primary trastuzumab-resistant and trastuzumab-sensitive cells, we performed Micro-C, a high-resolution chromatin conformation capture method, in JIMT1 and SKBR3 cells. Analysis of two replicates per cell line revealed strong concordance between replicates (Figure S2a–b). We observed differences in both intra- and interchromosomal contacts between the two cell lines. Intrachromosomal contacts involving chromosomes 15, 16, 17, and 21 were more abundant in JIMT1 cells than in SKBR3 cells, whereas those involving chromosomes 4, 8, and 14 were less abundant (Figure 2a). For interchromosomal contacts, interactions involving chromosomes 6, 14, and 22 were more prominent in SKBR3 cells, whereas those involving chromosomes 16 and 21 were stronger in JIMT1 cells (Figure 2b).

**Figure 2.**
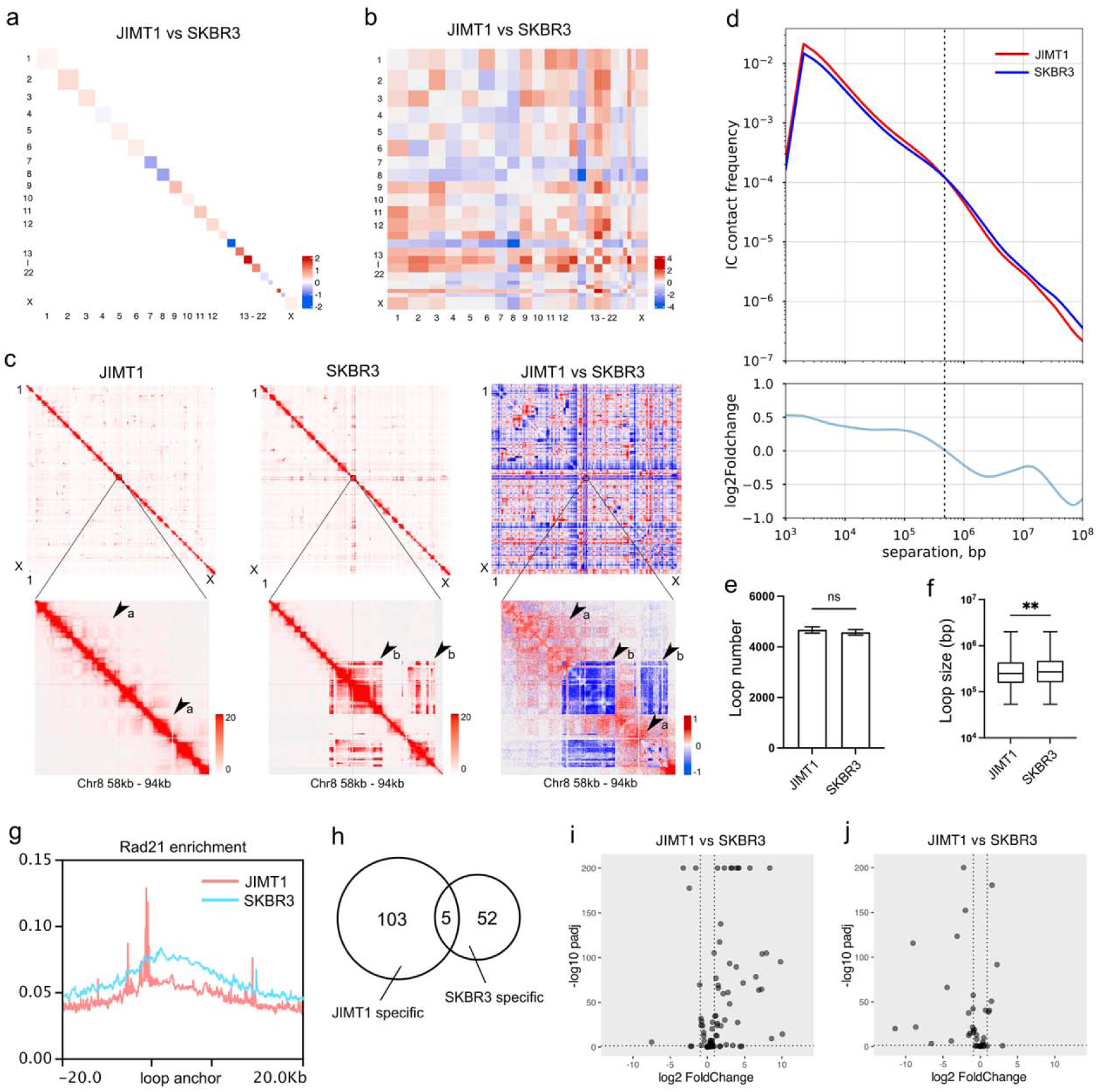
Differential chromatin architecture between trastuzumab-sensitive and resistant cells. **a**, Log2 fold changes of intra-chromosomal contacts (JIMT1 vs SKBR3). **b**, Log2 fold changes of inter-chromosomal contacts (JIMT1 vs SKBR3). **c**, Detailed intra-chromosomal contacts in a specific chromatin region (Chr8: 58 - 94 kb) of JIMT1 and SKBR3 cells (a indicates JIMT1-specific contacts, b indicates SKBR3-specific contacts). **d**, Relationship between contact frequency (P) of intra-chromosomal contacts and genome distance (s) in JIMT1 and SKBR3 cells (Top) and the log2 fold changes of frequency ranked by distance (JIMT1 vs SKBR3) (Bottom). The dotted line indicates the crossing point of two curves (∼240 kb). **e**, The number of chromatin loops in JIMT1 and SKBR3 cells. **f**, The distributions of loop length in JIMT1 and SKBR3 cells. **g**, The overall enrichment of the cohesin component Rad21 at the anchors of the chromatin loop in JIMT1 and SKBR3 cells. **h**, The number of distinct and shared promoter-enhancer chromatin loops in JIMT1 and SKBR3 cells. **i**, The expression of genes with JIMT1-specific promoter-enhancer chromatin loops. **j**, The expression of genes with SKBR3-specific promoter-enhancer chromatin loops.

Cell line-specific gains and losses of intrachromosomal contacts were also observed at individual genomic loci. For example, within the 58–94 kb region of chromosome 8, JIMT1-specific contacts were detected in the 58–70 kb and 81–86 kb intervals, whereas SKBR3-specific contacts were enriched in the remaining regions (Figure 2c). Analysis of contact frequency as a function of genomic distance using P(s) curves showed that JIMT1 cells had a higher frequency of intrachromosomal contacts at distances below approximately 240 kb. In contrast, longer-range contacts were more frequent in SKBR3 cells (Figure 2d).

Chromatin loops represent enriched contacts between pairs of genomic regions and can contribute to the regulation of gene expression. Although JIMT1 and SKBR3 cells exhibited similar numbers of loops (Figure 2e), loops were generally larger in SKBR3 cells (Figure 2f), consistent with the greater frequency of longer-range contacts in these cells. The genomic distribution of loop anchors was similar between the two cell lines, with most anchors located in promoters, introns, and exons (Figure S2c-d). Enrichment of RAD21, a component of the cohesin complex involved in maintaining chromatin loops, was higher at loop anchors in JIMT1 cells than in SKBR3 cells (Figure 2g).

Promoter–enhancer loops can bring regulatory elements into proximity with their target genes and facilitate transcriptional regulation. Although more than half of the identified loops connected two non-promoter regions, promoter–enhancer loops accounted for 2.14% of loops in JIMT1 cells and 1.88% in SKBR3 cells (Figure S2e-f). We identified 103 JIMT1-specific and 52 SKBR3-specific promoter–enhancer loops, together with five loops shared by both cell lines (Figure 2h). Most genes associated with cell line-specific promoter–enhancer loops showed higher expression in the corresponding cell line (Figure 2i-j). Genes associated with JIMT1-specific loops were primarily linked to pathways and processes implicated in treatment resistance, including p53 and TNF-α signaling, epithelial-mesenchymal transition, FoxO signaling, and adherens junctions (Figure S2g-h).

The activity-by-contact (ABC) score estimates the contribution of nearby enhancer activity and enhancer–promoter contacts to gene expression. The extended ABC-P^2^ score additionally incorporates promoter activity (Figure S2i). For genes associated with enhancers showing altered activity, we calculated and compared ABC scores between JIMT1 and SKBR3 cells, considering enhancers within ±1 Mb of each gene and their corresponding enhancer–promoter contact strengths. Differences in ABC scores were positively correlated with differences in transcription between the two cell lines (Figure S2j). Incorporating promoter activity into the ABC-P2 score strengthened this correlation (Figure S2k). Together, these findings suggest that promoter histone marks, enhancer activity, and enhancer–promoter contacts may act cooperatively to regulate gene transcription.

### Variant genomic compartment states link histone modifications

Principal component analysis (PCA) of Micro-C data was used to characterize chromatin compartments in JIMT1 and SKBR3 cells. This analysis identified differences in the distribution of A and B compartments, which are generally associated with active and repressive chromatin states, respectively.

At a resolution of 25 kb, approximately 34.6% of genomic bins exhibited stable compartment states between JIMT1 and SKBR3 cells. Opposite compartment assignments were observed in 9.45% of bins classified as A in JIMT1 and B in SKBR3 cells, and in 9.27% of bins classified as B in JIMT1 and A in SKBR3 cells. Additionally, 8.54% and 14.0% of bins exhibited stronger repressive compartment characteristics in JIMT1 and SKBR3 cells, respectively, whereas 12.4% and 9.27% exhibited stronger active compartment characteristics in the respective cell lines (Figure 3a).

**Figure 3.**
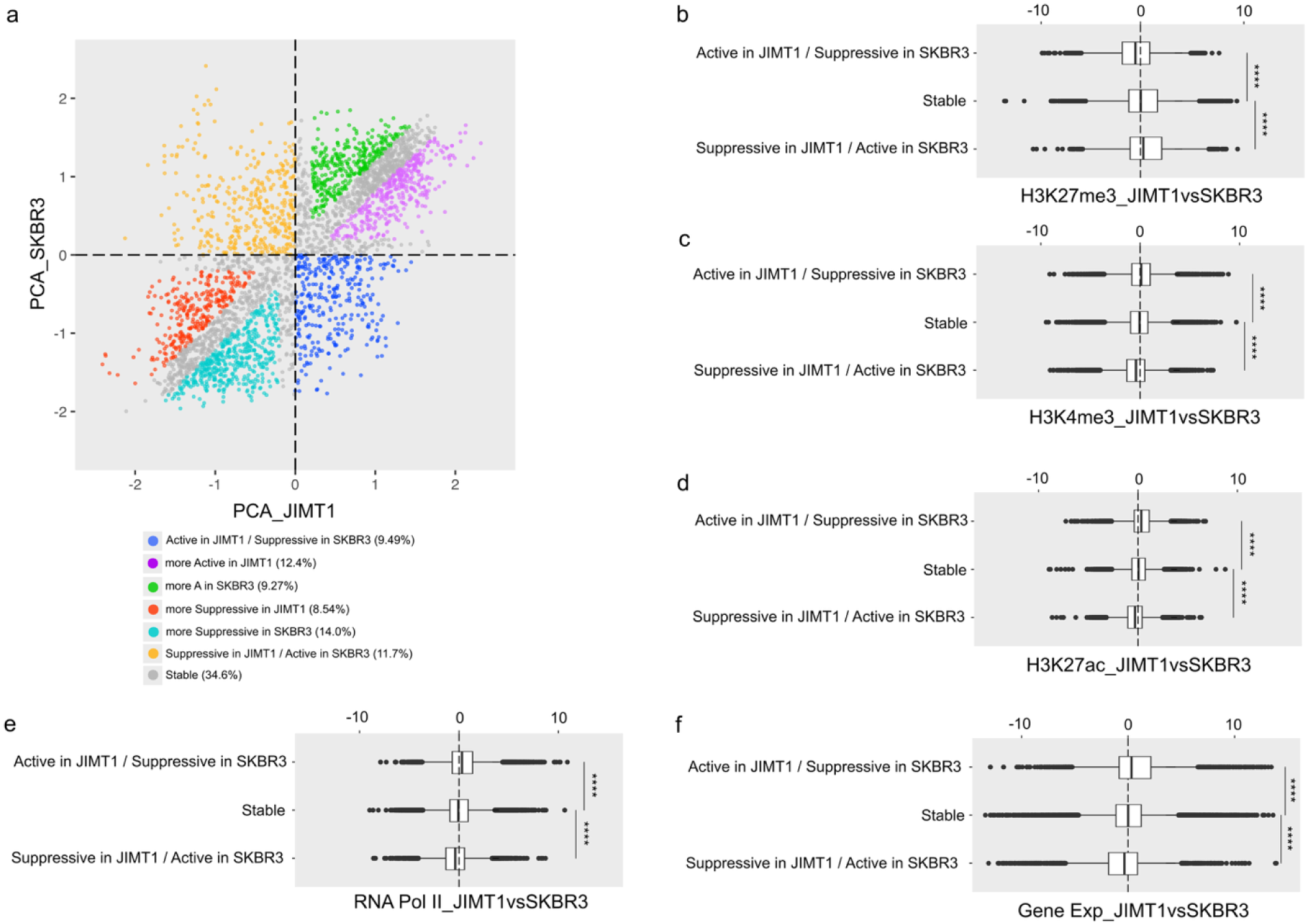
Differential A/B compartments between trastuzumab-sensitive and resistant cells. **a**, The distributions of compartment scores at 25 kb genomic bin resolution in JIMT1 and SKBR3 cells. Seven colors represent different categories of bins based on the types of their scores in the two cells (active in JIMT1 / suppressive in SKBR3, suppressive in JIMT1 / active in SKBR3, more active in JIMT1, more active in SKBR3, more suppressive in JIMT1, more suppressive in SKBR3 and stable). **b-f**, The distributions of log2 histone modifications(H3K27me3, H3K4me3 and H3K27ac), RNA polymerase and gene transcription changes in differential compartment score types in JIMT1 and SKBR3 cells (active in JIMT1 / suppressive in SKBR3, suppressive in JIMT1 / active in SKBR3 and stable).

Genome-wide distributions of histone modifications were associated with compartment states in both cell lines. The repressive mark H3K27me3 was enriched in B compartments relative to A compartments. In bins assigned to compartment A in JIMT1 cells and compartment B in SKBR3 cells, H3K27me3 enrichment was lower in JIMT1 than in SKBR3 cells. The opposite pattern was observed in bins assigned to compartment B in JIMT1 cells and compartment A in SKBR3 cells (Figure 3b). Similar trends were observed in bins with stronger active or repressive compartment characteristics in either cell line (Figure S3a).

Enrichment of the active histone marks H3K4me3 and H3K27ac, as well as RNA polymerase II, showed the opposite pattern to H3K27me3, with higher signals in A compartments than in B compartments. These signals were also enriched in bins exhibiting active compartment characteristics in either JIMT1 or SKBR3 cells (Figures 3c-e and S3b-d). Consistent with these patterns, genes located in A compartments showed higher transcriptional activity than those in B compartments in both cell lines (Figures 3f and S3e).

### Epigenetic alterations regulate crucial gene expression

By integrating gene expression data with cell type-specific promoter–enhancer interactions and promoter epigenomic profiles, we found that most genes associated with JIMT1-specific interactions exhibited higher promoter H3K4me3 enrichment, greater RNA polymerase II occupancy, and increased transcription in JIMT1 cells. A similar pattern was observed for genes associated with SKBR3-specific interactions in SKBR3 cells (Figure 4a).

**Figure 4.**
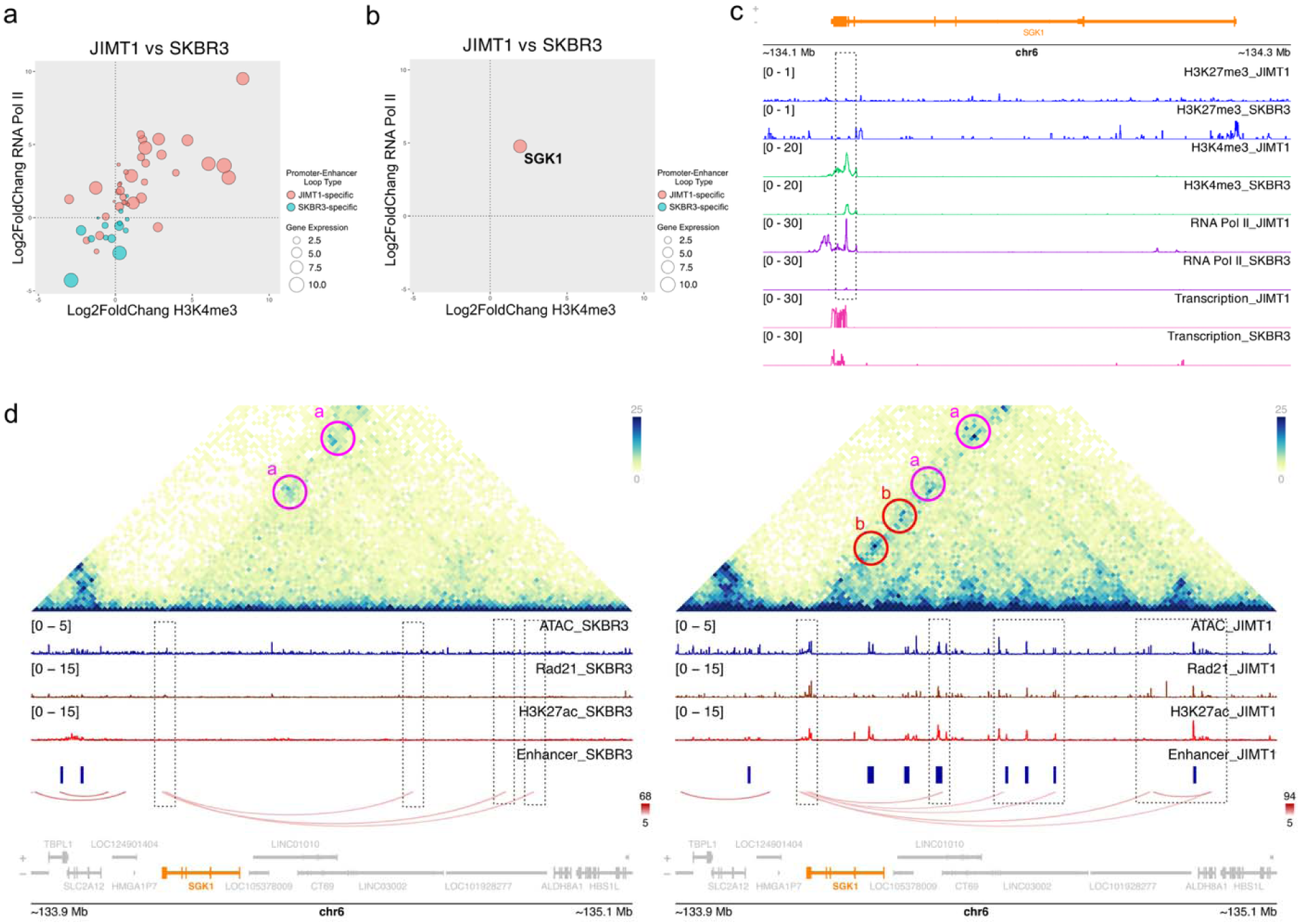
Altered histone modifications and DNA loops regulate gene transcription. **a**, The log2 fold changes of promoter H3K4me3 and RNA polymerase II, and transcription of genes with JIMT1 or SKBR3-specific promoter-enhancer chromatin loops. **b**, The log2 fold changes of promoter H3K4me3 and RNA polymerase II, and transcription of the SGK1 gene. **c**, The enrichments of promoter H3K27me3, H3K4me3 and RNA polymerase II, and transcription of SGK1 gene in JIMT1 and SKBR3 cells. **d**, The chromatin contacts, peaks of Rad21 and H3K27ac, activated enhancer and chromatin loops around SGK1 gene in JIMT1 and SKBR3 cells (red a represents chromatin contacts in both JIMT1 and SKBR3 cells, red b represents SKBR3-specific contacts).

Among the differentially expressed genes, SGK1 was a representative example of epigenetic activation, with elevated expression in JIMT1 cells (Figure 4b). SGK1 is associated with the mTOR, PI3K–AKT, and FoxO pathways and has been implicated in multiple cancer types. At the SGK1 promoter, H3K27me3 levels were similarly low in both cell lines, whereas H3K4me3 enrichment was approximately twofold higher in JIMT1 cells. This increase was accompanied by approximately fourfold higher RNA polymerase II occupancy and eightfold higher transcription (Figure 4c). These findings are consistent with promoter activation associated with an active histone mark rather than a reduction in the repressive H3K27me3 mark.

Changes in enhancer activity and chromatin architecture were also associated with SGK1 expression. In SKBR3 cells, the SGK1 promoter formed weak chromatin contacts, with low chromatin accessibility, low cohesin (RAD21) occupancy and limited interactions with active enhancers. In contrast, JIMT1 cells exhibited stronger and more frequent contacts connecting the SGK1 promoter to highly active enhancers, together with increased chromatin accessibility and cohesin enrichment (Figure 4d). Collectively, these observations suggest that coordinated differences in promoter histone modifications, enhancer activity, and three-dimensional chromatin organization contribute to SGK1 upregulation.

To investigate whether SGK1 is also involved in acquired trastuzumab resistance, we examined transcriptional and epigenomic data from a trastuzumab-resistant SKBR3 cell line established by our group (SKBR3_HR). SGK1 expression did not differ significantly between SKBR3 and SKBR3_HR cells (Figure S4a). Promoter enrichment of H3K27me3 and H3K4me3 at SGK1, as well as RNA polymerase II occupancy, was similar in SKBR3_HR and SKBR3 cells but lower than in JIMT1 cells (Figure S4b). Likewise, promoter–enhancer interactions, chromatin accessibility, and cohesin enrichment at the SGK1 locus were more similar between SKBR3_HR and SKBR3 cells than between SKBR3_HR and JIMT1 cells (Figure S4c). These findings indicate that SGK1-associated epigenomic changes differ between the intrinsic and acquired resistance models examined.

In contrast to SGK1, RNF152, which has been reported to exert tumor-suppressive effects in certain cancer types, exhibited SKBR3-specific promoter–enhancer contacts, chromatin accessibility, and cohesin enrichment (Figure S5a-d). Reduced promoter H3K4me3 enrichment and RNA polymerase II occupancy in JIMT1 cells were accompanied by lower RNF152 transcription than in SKBR3 cells. RNF152 expression was comparable in SKBR3 and SKBR3_HR cells and substantially higher in both than in JIMT1 cells. This expression pattern was accompanied by corresponding differences in enhancer activity, promoter–enhancer interactions, promoter H3K4me3 enrichment, and RNA polymerase II occupancy.

Together, the high SGK1 and low RNF152 expression observed in the model of primary trastuzumab resistance, but not in the model of acquired resistance, were accompanied by distinct epigenomic profiles. These findings suggest that SGK1 and RNF152 may be associated with primary trastuzumab resistance in HER2-positive breast cancer.

### SGK1 promotes malignant cellular phenotypes and is associated with trastuzumab resistance

To assess the functional contribution of SGK1 to trastuzumab resistance, we overexpressed SGK1 in SKBR3 cells and knocked it down in JIMT1 cells. Compared with control cells, SGK1-overexpressing SKBR3 cells exhibited increased colony formation, migration, and invasion (Figure 5a-d). Conversely, SGK1 knockdown in JIMT1 cells reduced proliferation, migration, and invasion (Figure S6a-d). SGK1 overexpression was also associated with increased phosphorylation of ERK and mTOR (Figure 5e-g), whereas SGK1 knockdown reduced the phosphorylation of both proteins (Figure S6e-f). These findings implicate SGK1 in signaling pathways associated with cell survival and proliferation. Furthermore, tumor tissues from patients with trastuzumab-resistant disease exhibited higher SGK1 enrichment than those from patients with trastuzumab-sensitive disease (Figure 5h), supporting an association between SGK1 and trastuzumab resistance.

**Figure 5.**
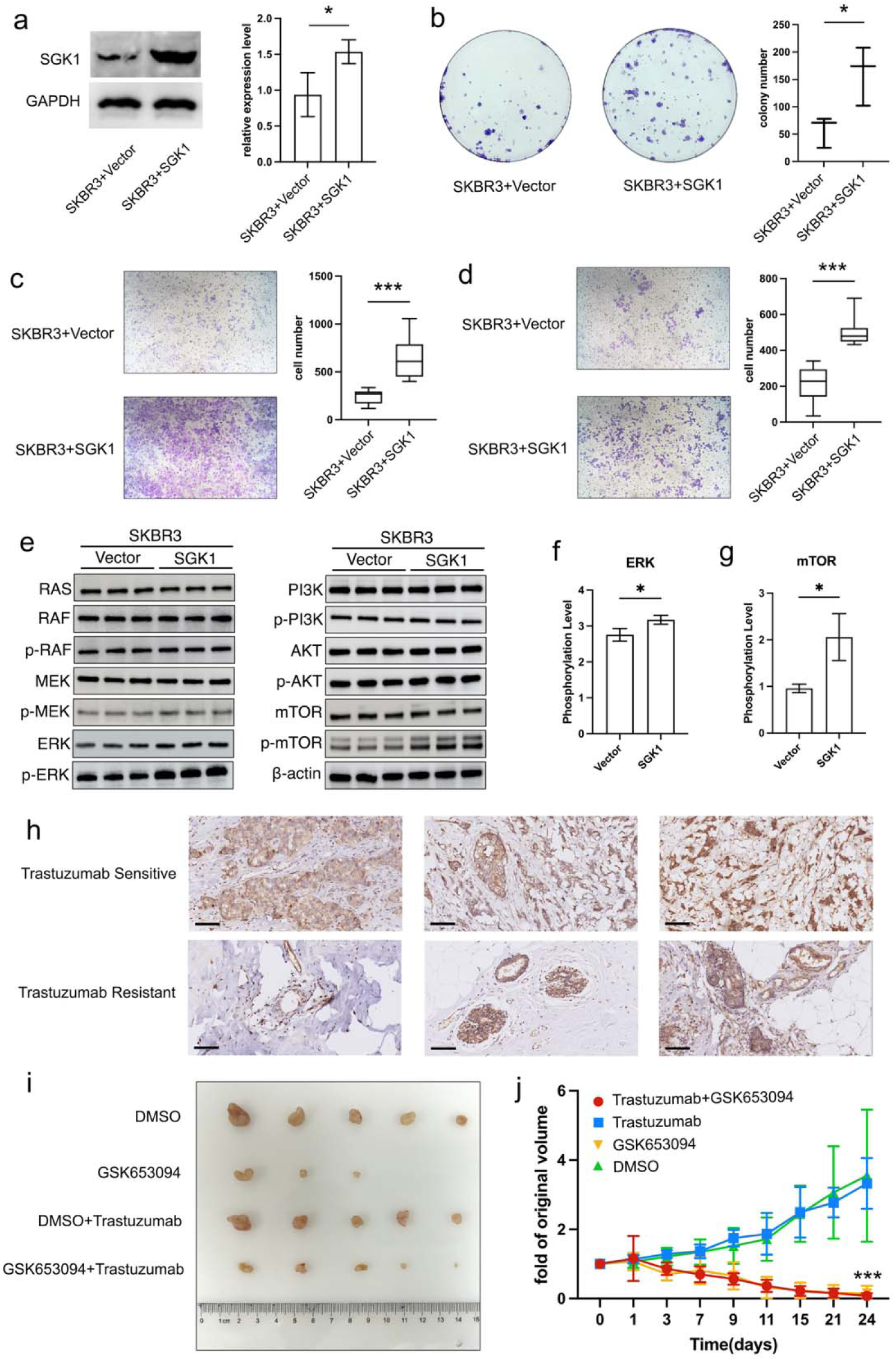
Upregulated SGK1 expression connects to trastuzumab resistance. **a**, The SGK1 protein level in SGK1- and vector-transfected SKBR3 cells. **b**, The clone formation ability of SGK1-overexpressing and normal SKBR3 cells. **c**, The migration ability of SGK1-overexpressing and normal SKBR3 cells. **d**, The invasive abilities of SGK1-overexpressing and normal SKBR3 cells. **e**, The expression and phosphorylation levels of major proteins in mTOR and ERK pathways in SGK1-overexpressing and normal SKBR3 cells. **f**, The phosphorylation levels of ERK in SGK1-overexpressing and normal SKBR3 cells. **g**, The phosphorylation levels of mTOR in SGK1-overexpressing and normal SKBR3 cells. **h**, SGK1 levels in the breast cancer tissues of trastuzumab-sensitive and resistant patients (scale bar: 100 μm). **i**, Tumor samples of different treatment strategies. **j**, Tumor volume changes of different treatment strategies during the experiment.

To evaluate the therapeutic potential of targeting epigenetically activated pathways, we tested the SGK1 inhibitor GSK653094 alone and in combination with trastuzumab in a JIMT1 xenograft mouse model. Tumor growth in the trastuzumab-only group was comparable to that in the untreated control group, with tumor volumes reaching nearly three times their initial values. In contrast, GSK653094 treatment, either alone or in combination with trastuzumab, markedly suppressed tumor growth (Figure 5i-j).

## Discussion

Genetic and epigenetic alterations are critical drivers of cancer progression, operating through distinct yet interrelated mechanisms. While genetic alterations—such as mutations, gene amplifications, and chromosomal rearrangements—permanently modify DNA sequences or gene copy numbers, epigenetic modifications are typically reversible and dynamic. These epigenetic changes, including histone modifications and chromatin architecture alterations, regulate transcriptional activity without altering the underlying genetic code, thereby influencing gene expression patterns associated with tumor development and progression.

Genetic alterations are generally regarded as the primary foundation of intrinsic drug resistance. For instance, HER2-positive breast cancer is characterized by the amplification of the ERBB2 gene, while mutations in ERBB2 or PIK3CA can contribute to resistance to trastuzumab^16,17^. In addition, intrinsic epigenetic modifications can also lead to alternative gene expression, contributing to cellular variability and potentially influencing drug resistance. DNA methylation is the most extensively studied epigenetic modification in the context of carcinogenesis and tumor progression^18–20^. The silencing of several tumor suppressor genes, including BRCA1, TP53, and CDH1, is commonly mediated by hypermethylation of CpG islands within their promoter regions^21–24^.

Differential modifications of histones can profoundly influence gene expression and drive tumor progression. The increased deposition of the repressive histone mark H3K27me3 at the promoter regions of tumor suppressor genes, such as BRCA1 and TP53, can lead to transcriptional silencing, thereby facilitating tumor progression^25–27^. In contrast, elevated levels of the activating histone mark H3K4me3 at the promoters of oncogenes, such as MYC, can significantly enhance their transcriptional activity, promoting carcinogenesis^28–31^. Furthermore, the overexpression of several oncogenes in breast cancer may be associated with an increased level of the enhancer mark H3K27ac^32–34^.

Furthermore, chromatin structure alteration also plays a crucial role in carcinogenesis and tumor progression and should not be overlooked. On one hand, chromatin accessibility, determined by nucleosome positioning, facilitates the binding of transcription factors or RNA polymerases to enhancer and promoter regions^35^. For instance, in estrogen receptor-positive (ER-positive) breast cancers, the binding of the activated estrogen receptor (ERα) is associated with changes in chromatin accessibility^36,37^. On the other hand, alterations in three-dimensional genomic architecture, particularly chromatin loops, can modulate the interactions between promoters of oncogenes or tumor suppressor genes and their nearby or distant enhancers, influencing their transcriptional activity and tumor progression^38–40^. In ER-positive breast cancers, the binding of ERα can promote chromatin looping between enhancers and ERα-responsive genes in cooperation with FOXA1 and GATA3^41–44^. Similarly, in triple-negative breast cancer (TNBC), the aberrant expression of MYC, BRD4, and SOX9 can be driven by chromatin loops between their promoters and associated superenhancers^45–47^.

In this study, we identified several notable epigenetic differences at promoter and enhancer regions. Among these, the change in the activating histone mark H3K4me3 at the promoter exhibited a stronger correlation with gene transcription than the repressive mark H3K27me3, suggesting that H3K4me3 may play a more prominent role in gene expression regulation. This finding bears a resemblance to our previous observations in acquired trastuzumab resistance^48^. Additionally, the importance of chromatin interactions and the reprogramming of enhancer activity, marked by alterations in non-promoter H3K27ac peaks, was evident in both intrinsic and acquired trastuzumab resistance^48^. The binding of transcription factors to these activated enhancer regions would be enhanced, promoting RNA polymerase II recruitment, facilitated by the formation of chromatin loops connecting single or multiple enhancers to specific promoters.

Combining data on altered histone modifications and promoter-enhancer interactions, we identified several genes exhibiting strong correlations between their transcriptional activity and the dynamics of their promoters or nearby enhancers. In this study, we primarily focused on genes with a robust positive correlation between these two factors. Of them, we suggested the SGK1 gene, which emerged as particularly significant, playing a crucial role in cancer-related cellular processes. As a serine/threonine kinase, SGK1 is known to be involved in a variety of critical cellular signaling and metabolic pathways. Dysregulation of SGK1 has been closely associated with several severe diseases, including cardiovascular, neurological, and metabolic disorders, as well as colorectal, lung, and breast cancers^26,49–54^. The classical function of SGK1 encompasses the activation of Ca²⁺ and K⁺ channels, along with amino acid and glucose transporters^26,49^. Additionally, its involvement in the PI3K-Akt, mTOR, and FoxO3 pathways is well-documented^55–58^. A few studies have highlighted the potential of SGK1 as a prognostic marker in breast cancer, suggesting its significance in predicting invasive disease^54,59,60^. Furthermore, SGK1 inhibition has been shown to disrupt cancer metastasis^55,61^.

In our study, SGK1 overexpression increased colony formation, migration, and invasion, while changes in SGK1 expression were also accompanied by changes in ERK and mTOR phosphorylation. These results support a role for SGK1 in the cellular phenotypes and signaling readouts measured and could be a potential mechanism to stimulate survival and proliferation and contribute to intrinsic trastuzumab resistance in certain cell models, but extending this to a universal and complete signaling mechanism still needs further experiments.

In summary, our study highlights the epigenetic differences, including histone modifications (H3K27me3, H3K4me3, and H3K27ac) and alterations in genomic architecture, between intrinsic trastuzumab-resistant and sensitive cells. Additionally, we propose that the promoter-associated H3K4me3, rather than H3K27me3, in conjunction with activated enhancers and rearranged promoter-enhancer interactions, plays a coordinated role in regulating gene transcription. By integrating these data, we suggest that the increased transcription of the SGK1 gene in trastuzumab-resistant cells may be driven by the enhanced presence of promoter H3K4me3, activated nearby enhancers, and reinforced promoter-enhancer contacts. Targeting SGK1 activity may significantly restrict tumor cell function in both *in vitro* and *in vivo* models, offering a potential therapeutic strategy for treating intrinsic trastuzumab-resistant HER2-positive breast cancer in clinical settings.

## Materials and methods

### Cell culture and transfection

Primary HER2-positive trastuzumab-sensitive breast cancer SKBR3 cells and trastuzumab-resistant breast cancer JIMT1 cells were cultured in modified Eagle’s medium (DMEM) (Gibco, 11960044) supplemented with 10% fetal bovine serum (FBS) (Gibco, 16140071), 80 U/ml penicillin, and 0.08mg/ml streptomycin (Gibco, 15140122) at 37 ℃ in a humidified environment with 5% CO_2_. The acquired trastuzumab-resistant SKBR3_HR cells, which were established via long-term and gentle gradient trastuzumab treatment from 1ug/ml to 20ug/ml in 30 weeks in our previous study^48^, were also cultured in DMEM supplemented with 10% FBS.

Transient transfection of plasmid DNA or siRNA was performed with Lipofectamine 3000 (Thermo, L3000001) according to the manufacturer’s protocol. Plasmid DNA or siRNA with P3000 was diluted in Opti-MEM, mixed with the same diluted Lipofectamine 3000, and then added to cells. After 48 hours of incubation, treated cells were harvested for the following experiments.

### Cell viability assay

Cell viability was measured with a CCK-8 kit (Vazyme, A311). 5000 cells were seeded in a 96-well plate. After 24 h incubation, the old medium was replaced by the fresh drug-containing medium. After another 48 h culture, the fresh CCK-8 containing medium was applied to replace the old medium for another 1 h incubation, the absorbance of each well at 450 nm wavelength was measured by a multi-well plate reader.

### Cell proliferation assay

Cell proliferation was assessed using a colony formation assay. Cells were harvested, counted, and seeded at a density of 500 cells per well in 6-well plates. The plates were incubated at 37°C in a humidified environment with 5% CO₂ for one week. Following incubation, cells were rinsed with PBS, fixed with 4% formaldehyde, and stained with 0.5% crystal violet for 30 minutes at room temperature. The stained cells were then gently rinsed with water and allowed to dry completely. Colonies were captured with a microscope and counted using ImageJ.

### Cell migration assay

Cell migration was assessed using a transwell assay. Briefly, 1 × 10⁵ cells were seeded into the upper chamber and incubated at 37°C with 5% CO₂ for 24 hours. Following incubation, cells on the lower surface of the membrane were fixed with 4% formaldehyde, stained with 0.5% crystal violet, and washed with PBS. Migrated colonies were captured and then quantified using ImageJ.

### Cell invasion assay

Cell migration was assessed using a transwell assay with a Matrigel-coated membrane in the upper chamber. The membrane was first coated with serum-free medium containing diluted Matrigel and incubated at 37°C for 2 hours. Subsequently, 1 × 10⁵ cells were seeded into the upper chamber and incubated at 37°C with 5% CO₂ for 24 hours. After incubation, cells on the lower surface of the membrane were fixed with 4% formaldehyde, stained with 0.5% crystal violet, and washed with PBS. Migrated colonies were captured and then quantified using ImageJ.

### Animal study

JLMT1 cells were injected into the lower flanks of 5-week-old BALB/c nude mice. When the tumor volume reached approximately 40 mm³, the mice were treated with trastuzumab (10 mg/kg) or DMSO on the 1st, 3rd, and 5th days of each week, and GSK-653094 (150 µg/day) or DMSO from the 1st to the 5th day of each week for 3 weeks via intraperitoneal injection. Tumor volume was calculated using the formula π/6 × width² × length.

### Protein extraction and western blot

Cells were washed with PBS after collection and lysed in an SDS lysis solution(Beyotime, P0013G) supplied with a protein inhibitor cocktail (MCE, HY-K0011) on ice for 5 min. Lysates were centrifuged at 12000 rpm for 10 min at 4 ℃. The supernatants were mixed with 5X loading buffer (Beyotime, P0015) and boiled at 95 ℃ for 15min. Proteins were separated by SDS-PAGE with 15% gels, transferred to PVDF membranes (Bio-Rad, 1620177) and blocked in 5% skim milk in TBST (20mM Tris, 150mM NaCl, and 0.1% Tween-20) for 1 h. Membranes were incubated in TBST with primary antibodies (SGK1 (Proteintech, 28454-1-AP)) at 4 ℃ overnight. After being washed with TBST 3 times, membranes were incubated in TBST with horseradish peroxidase (HRP)- conjugated anti-rabbit secondary antibody at room temperature for 1 h. After being washed with TBST 3 times, protein bands were visualized in a Western Blot imaging system with an ECL chemiluminescence kit (Vazyme, E422-01) and quantified using ImageJ.Sequencing library preparation and data analysis for the CUT&Tag assay

Cleavage under targets and tagmentation (CUT&Tag) sequencing was applied to identify genomic histone modifications and the binding of RNA polymerase II and Rad21 with a commercial CUT&Tag assay kit (Vazyme, TD903). In brief, 10^5^ cells with > 90% viability were obtained for each test. After binding with magnetic Concanavalin A beads, cells were permeabilized and treated with primary antibodies (H3K27me3(CST, #9733), H3K4me3(Abcam, ab213224), H3K27Ac (Abcam, ab177178), RNA pol II CTD (Abcam, ab300575) and Rad21 (Abcam, ab217678)) and the following goat anti-rabbit secondary antibody separately. After that, the genomic DNA of each sample was first digested by protein A/G bonded Tnp, then released by proteinase K treatment and purified with SPRIselect beads (Vazyme, N411). Library amplification was performed by an Illumina-compatible index kit (Vazyme, TD202), and the following sequencing was carried out on an Illumina NovaSeq 6000 platform.

Raw paired CUT&Tag sequencing reads were aligned to the human hg38 reference genome with Bowtie2 (v2.2.5) and then filtered and converted to .bed files by Samtools (v1.14) and Bedtools (v2.30.0). SEACR (v1.3) was applied for peak calling, while DEseq2 (v1.44.0) and chromVAR (v1.26.0) packages in R were used for differential analysis and annotation. Significantly changed peaks were defined as Padj <= 0.05 & |log2 fold change| >= 1.

### Sequencing library preparation and data analysis for the Micro-C assay

The Micro-C sequencing libraries were prepared using a commercial Micro-C Kit (Dovetail #21006) according to the manufacturer’s protocol. Briefly, the chromatin was fixed and cross-linked with disuccinimidyl glutarate (DSG) and formaldehyde in the nucleus and then digested in situ with micrococcal nuclease (MNase). After that, chromatin fragments were released from cells by SDS treatment and bound to Chromatin Capture Beads. Next, the chromatin ends were repaired and ligated to a biotinylated bridge adapter, followed by proximity ligation of adapter-containing ends. After proximity ligation, the crosslinks were reversed, the associated proteins were degraded, and the DNA was purified and then converted into a sequencing library using Illumina-compatible adaptors. Biotin-containing fragments were isolated using streptavidin beads before PCR amplification. The sequencing was also performed with an Illumina NovaSeq 6000 platform.

Raw paired Micro-C sequencing reads were aligned to the human hg38 reference genome assembly with BWA-MEM (v0.7.17). The following ligation junction finding, sorting and PCR duplicate removal were all performed with pairtools (v1.0.2). Juicertools (v1.22.1) and cooler (v0.9.2) were applied to generate .hic and .mcool contact matrices. The stratum-adjusted correlation coefficient (SCC), indicating the concordance between biological replicates of each cell type, was calculated by the hicrep (v1.12.2) package in R. The P-s curve was generated using cooltools (v0.5.4) as a function of intra-chromosome contact frequency (P) and genomic distance (s). HOMER (V4.11) was used to identify TADs and loops on contact matrices. ABC scores were calculated by combining enhancer activity and enhancer–promoter contact strength, as described previously^62^. For each gene, active enhancers were defined as significant H3K27ac peaks within ±1 Mb of the promoter. Their normalized H3K27ac signals were used as enhancer activity scores (*A*), and normalized contact strengths at 5-kb resolution were used as contact scores (*C*). The ABC score was calculated as the sum of *A* × *C* across all candidate enhancers. Promoter activity (*P*) was defined as the calibrated H3K4me3 signal within ±2.5 kb of the transcription start site. The ABC-P2 score was calculated as ABC × *P*².

Whole-genome compartment analyses (PCA) on the Micro-C contact matrix at 25kb resolution within 100kb windows and the following PC1-based compartment discovery were performed by HOMER. Different compartment changes were defined similarly as previous studies: A to stronger A (JIMT1 PC1 – SKBR3 PC1 > 0.2 & JIMT1 PC1 > 0.2), B to A (JIMT1 PC1 > 0 & SKBR3 PC1 < 0), B to weaker B (JIMT1 PC1 – SKBR3 PC1 > 0.2 & JIMT1 PC1 < −0.2), B to stronger B (JIMT1 PC1 – SKBR3 PC1 < −0.2 & SKBR3 PC1 < −0.2), A to B(JIMT1 PC1 < 0 & SKBR3 PC1 > 0), A to weaker A (JIMT1 PC1 - SKBR3 PC1 < −0.2 & SKBR3 PC1 > 0.2), and the rest were all considered as stable.

## Statistical analysis

Statistical analyses were performed with GraphPad Prism and R. Student’s t-test was applied for two-group comparison. P < 0.05 was considered the significance limit and marked as * (P < 0.05), ** (P < 0.01), *** (P < 0.001), and **** (P < 0.0001). Three independent biological replicates were used for the CUT&Tag assay, while two independent biological replicates were used for the Micro-C assay.

## Acknowledgment

We thank Dr. Xiang Huang, Dr. Yan Liang, Dr. Fan Yang, Dr. Yanting Sun, Dr. Chunxiao Sun, Dr. Huizhi Gong, and Shuang Hu for helpful discussions. This project was supported by the National Natural Science Foundation of China (82573066, 82272667, and 81972484 to Yongmei Yin and 82203488 to Ningjun Duan), the Major Basic Research Fund of Jiangsu Province Hospital (QY202406), Clinical Diagnosis and Treatment Technology Innovation “Open Bidding and Leader Selecting” Project of Jiangsu Province Hospital (CZ1420240207), Special Program for Top-notch Innovative Talents of The First Affiliated Hospital with Nanjing Medical University (NJMUTY20250070), and the Collaborative Innovation Center for Tumor Individualization Program (JX21817902/008)

## Author contributions

NJ. D. conceived the project. NJ. D., YJ. H., ZX. Z. and N. J performed the wet lab experiments. NJ. D. and YJ. H. conducted dry lab work. NJ. D, YJ. H. and ZX. Z. wrote the manuscript and arranged the figures. All authors reviewed the manuscript.

## Data availability

RNA-Seq data were deposited in GEO (accession number GSE297774). CUT & Tag data were deposited in GEO (accession number GSE297715)

## Supplementary Figures

**Figure S1.**
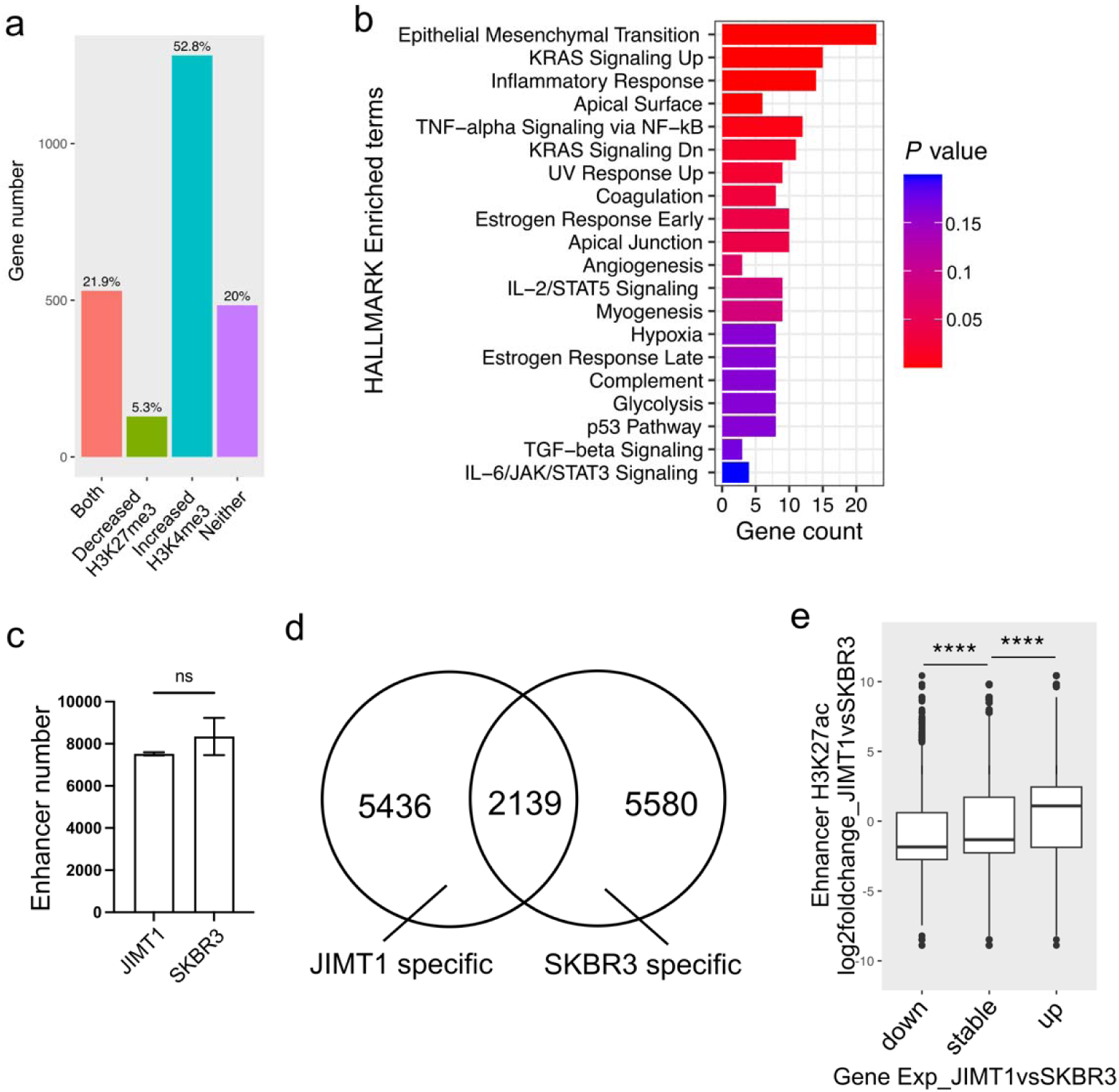
**a**, H3K27me3 and H3K4me3 changes at the promoter regions of upregulated genes in JIMT1 vs SKBR3 cells. **b**, GO HALLMARK enrichment of genes with activated bivalent promoters in JIMT1 cells. **c**, The total number of enhancers in JIMT1 and SKBR3 cells. **d**, The number of distinct and shared enhancers in JIMT1 and SKBR3 cells.

**Figure S2.**
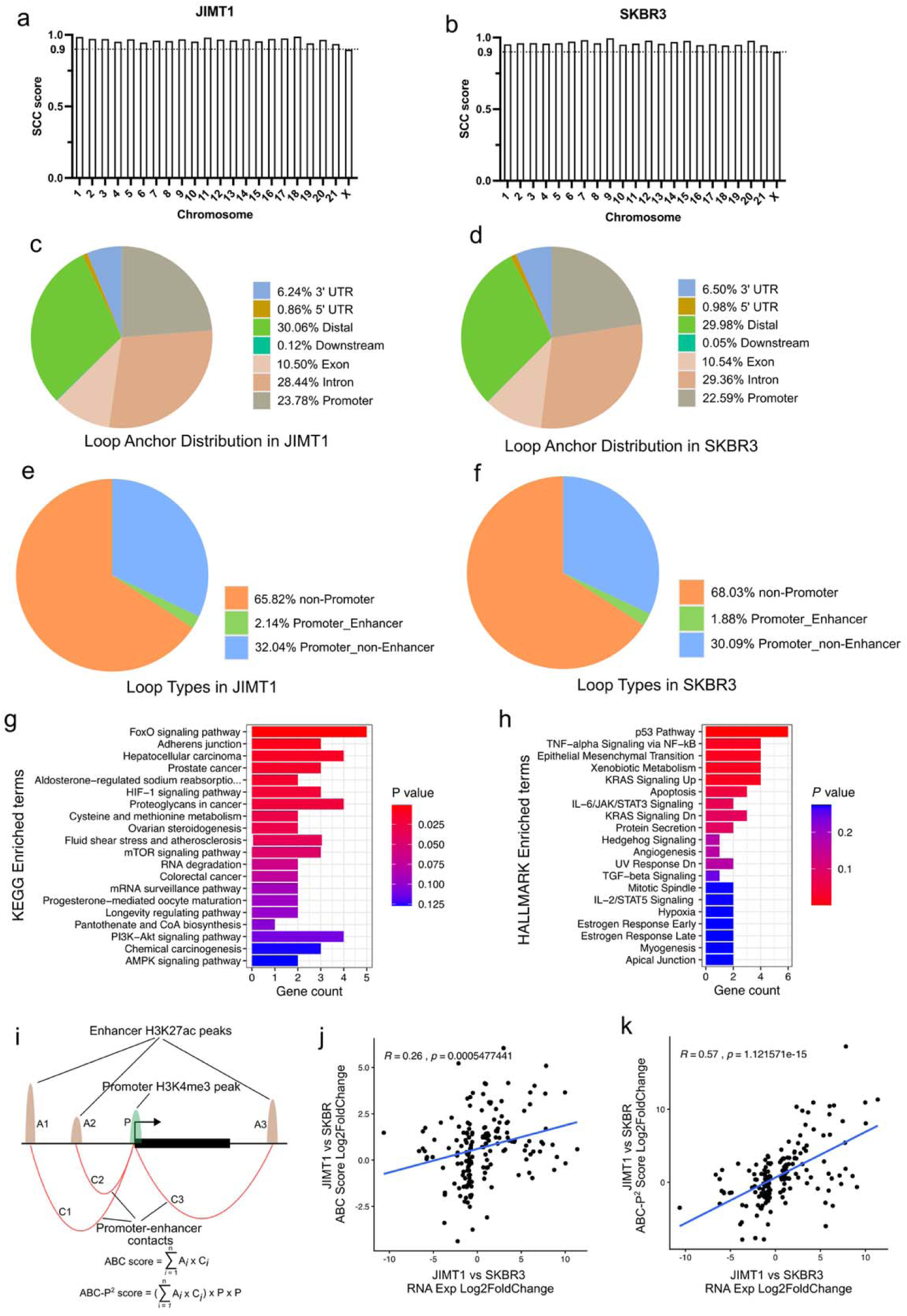
**a,** SCC correlation coefficients of different chromosomes between two replicates of JIMT1 cells. **b**, SCC correlation coefficients of different chromosomes between two replicates of SKBR3 cells. **c**, The distributions of chromatin loop anchors in JIMT1 cells. **d**, The distributions of chromatin loop anchors in SKBR3 cells. **e**, The types of chromatin loops in JIMT1 cells. **f**, The types of chromatin loops in SKBR3 cells. **g**, GO KEGG enrichment of genes with JIMT1-specific promoter-enhancer chromatin loops. **h**, GO HALLMARK enrichment of genes with JIMT1-specific promoter-enhancer chromatin loops. i, ABC model indicates that gene expression is affected by enhancer activity and promoter – enhancer contacts, while ABC-P^2^ additionally considers the strength of promoter activity. **j**, The Pearson correlation between the log2 fold changes of ABC score and RNA expression across JIMT1 and SKBR3 cells. **k**, The Pearson correlation between the log2 fold changes of ABC-P^2^ score and RNA expression across JIMT1 and SKBR3 cells.

**Figure S3.**
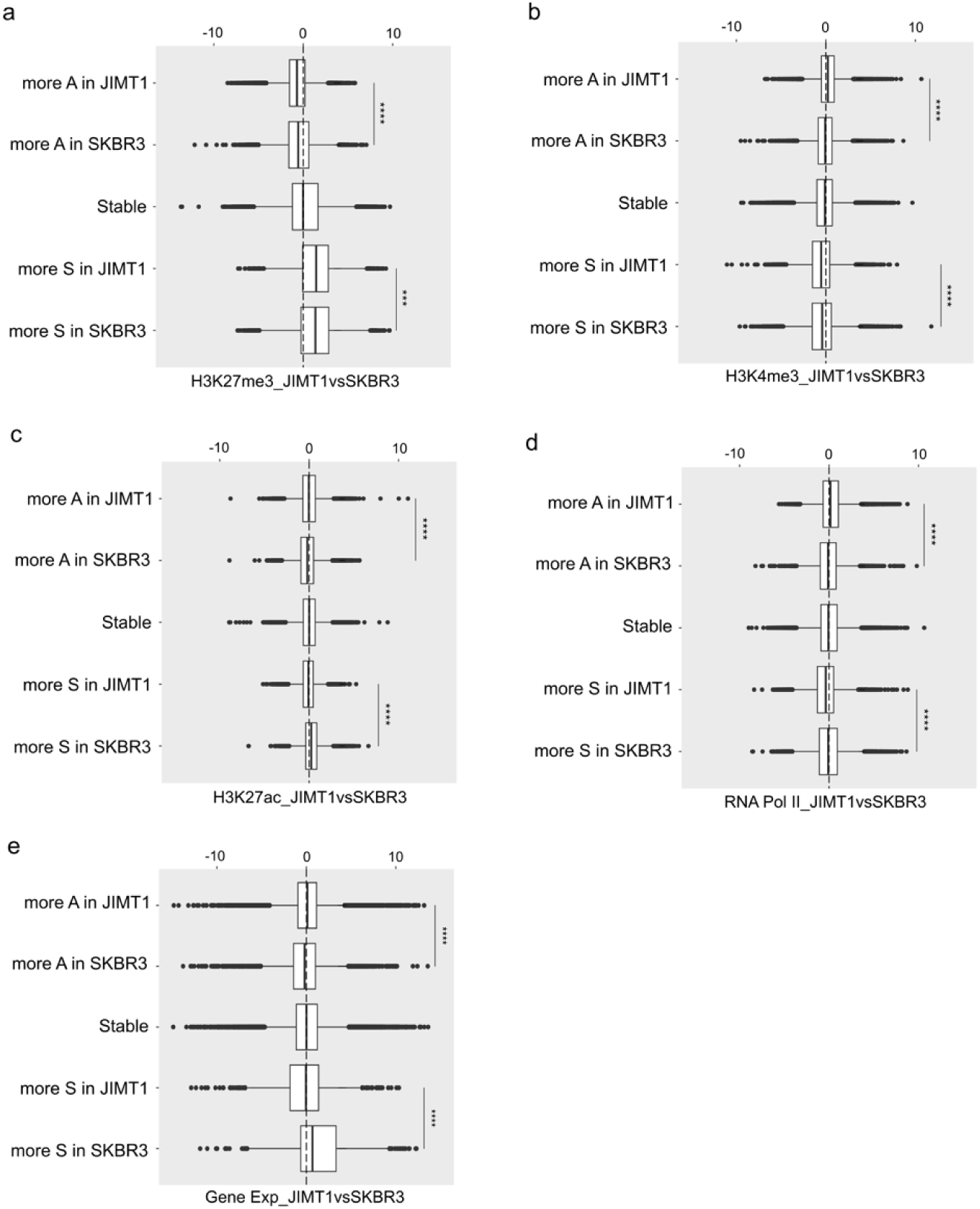
**a-e**, The distributions of log2 histone modifications(H3K27me3, H3K4me3 and H3K27ac), RNA polymerase and gene transcription changes in differential compartment score types in JIMT1 and SKBR3 cells (more active in JIMT1, more active in SKBR3, more suppressive in JIMT1, more suppressive in SKBR3 and stable).

**Figure S4.**
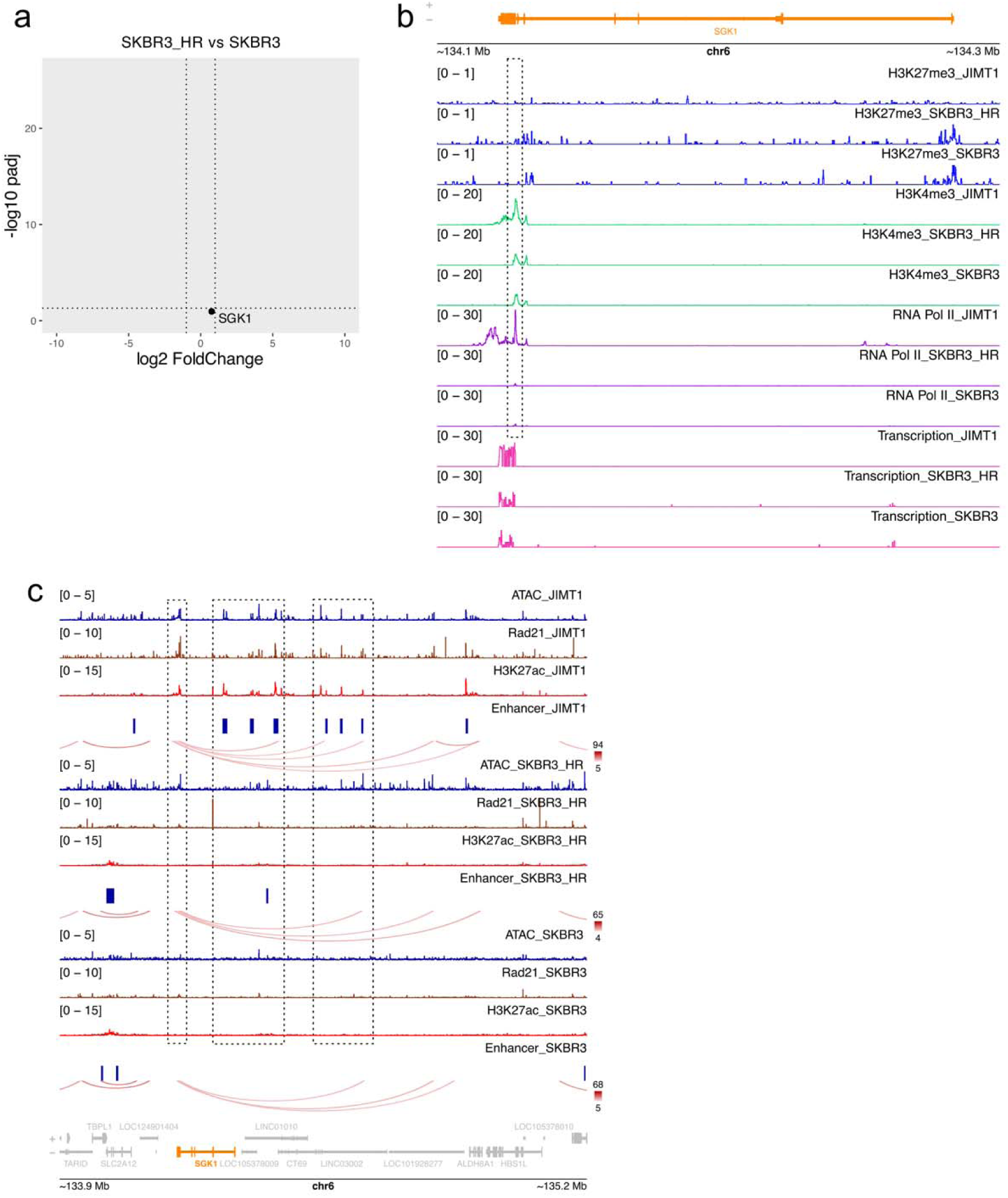
**a,** The expression of SGK1 gene in SKBR3_HR and SKBR3 cells. **b**, The levels of promoter H3K27me3, H3K4me3 and RNA polymerase II, and transcription of SGK1 gene in JIMT1, SKBR3 and SKBR3_HR cells. **c**, The peaks of Rad21 and H3K27ac, activated enhancer and chromatin loops around SGK1 gene in JIMT1, SKBR3 and SKBR3_HR cells.

**Figure S5.**
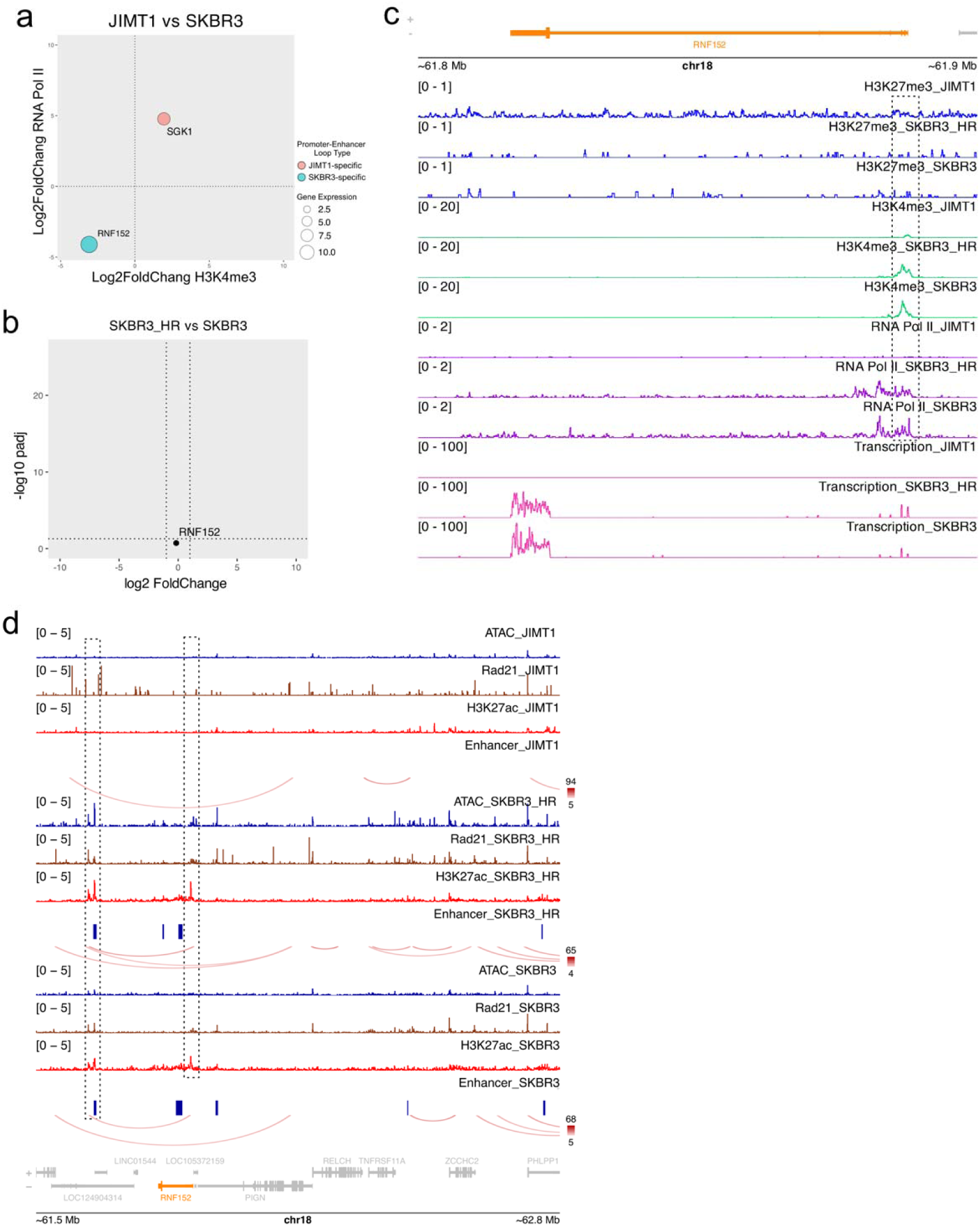
**a,** The log2 fold changes of promoter H3K4me3 and RNA polymerase II, and transcription of the RNF152 gene. **b**, The expression of RNF152 gene in SKBR3_HR and SKBR3 cells. **c**, The levels of promoter H3K27me3, H3K4me3 and RNA polymerase II, and transcription of RNF152 gene in JIMT1, SKBR3 and SKBR3_HR cells. **d**, The peaks of Rad21 and H3K27ac, activated enhancer and chromatin loops around RNF152 gene in JIMT1, SKBR3 and SKBR3_HR cells.

**Figure S6.**
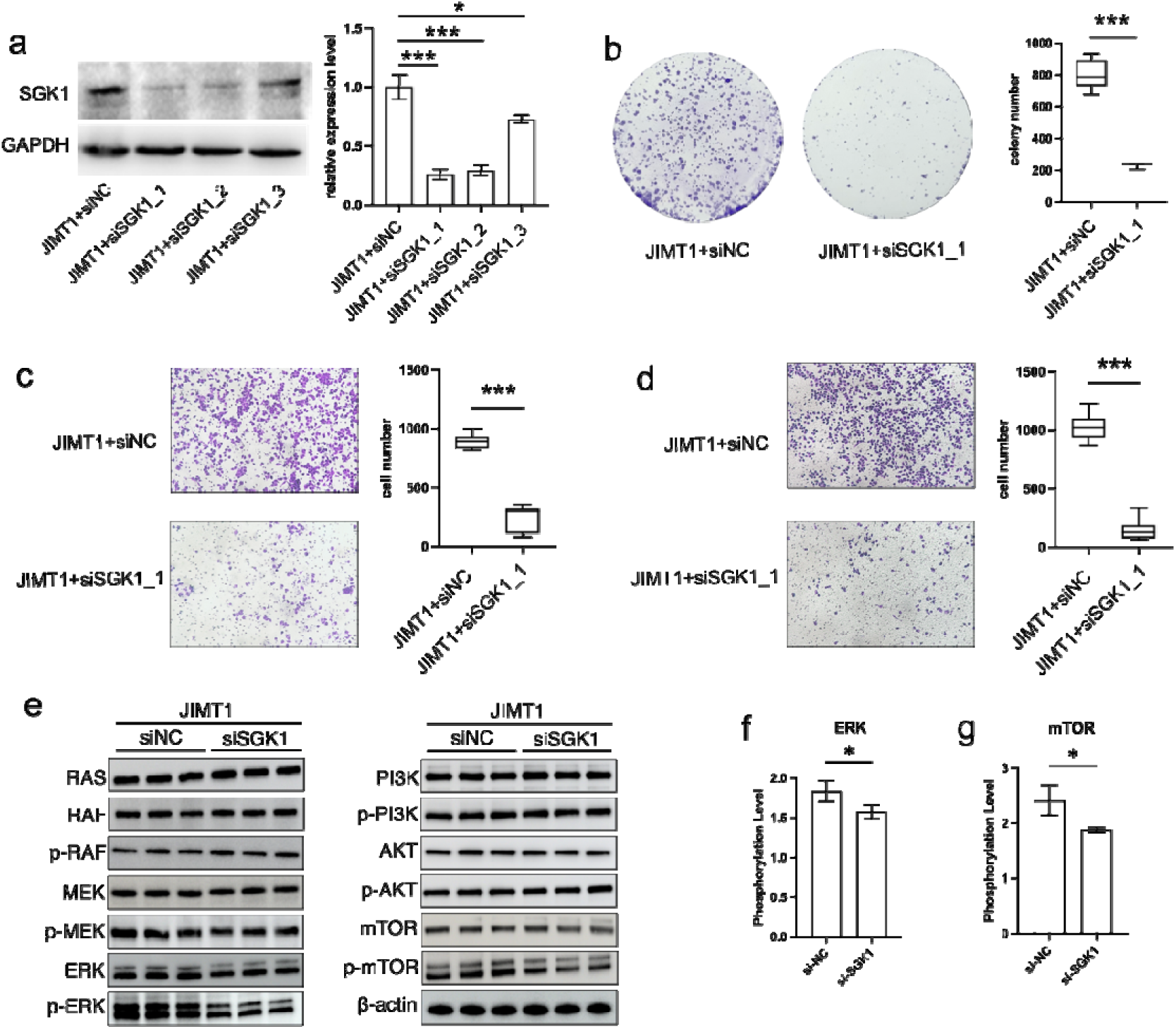
**a,** The SGK1 protein level in SGK1-knockdown and normal JIMT1 cells. **b**, The clonogenic ability of SGK1-knockdown and normal JIMT1 cells. **c**, The migration abilities of SGK1-knockdown and normal JIMT1 cells. **d**, The invasive abilities of SGK1-knockdown and normal JIMT1 cells. **e**, The expression and phosphorylation levels of major proteins in mTOR and ERK pathways in SGK1-knockdown and normal JIMT1 cells. **f**, The phosphorylation levels of ERK in SGK1-knockdown and normal JIMT1 cells. **g**, The phosphorylation levels of mTOR in SGK1-knockdown and normal JIMT1 cells.

